# Current estimates of population resilience are not robust to the longer-term impacts of directional environmental shifts

**DOI:** 10.1101/2025.04.04.647258

**Authors:** James Cant, Christina M. Hernández, Rachael H. Thornley, Andrew Hector, Iain Stott, Roberto Salguero-Gómez

## Abstract

Natural systems worldwide are exposed to gradual changes imposed by directional environmental shifts, known as ramp disturbances. However, contemporary assessments of population resilience focus on the resistance and recovery of systems following one-off (*i.e.*, pulse) disturbances. Thus, our current perception of demographic resilience overlooks the potential impacts of the cumulative effects of ramp disturbances on population dynamics. Simulating 50-year ramp disturbance scenarios upon the survival and fecundity dynamics of 511 populations across 344 species, we illustrate non-linear patterns in how directional environmental shifts will reshape the resilience of natural populations. Accordingly, existing estimates of population resilience are not robust against a backdrop of ongoing and continuous environmental change. Instead, we demonstrate the need to quantify how the relative partitioning of energetic resources across survival *vs.* fecundity informs population resilience. Our findings challenge the use of well-established approaches in forecasting population resilience to ongoing global change.

## Introduction

The urgent need to predict the responses of ecological systems to ongoing global change has fueled decades of research into the demographic resilience of ecological systems, that is, their ability to resist and recover from disturbances (Capdevila *et al*. 2021; Holling 1973, 1996). Common to our understanding of demographic resilience, however, is a focus on the implications of pulse disturbances (Van Meerbeek *et al*. 2021). Pulse disturbances impose an abrupt and discrete impact on some parameter(s) of the ecological system under consideration (Bender *et al*. 1984). Yet in reality, complex disturbance regimes characterised by gradual changes imposed by directional environmental shifts, or *ramp disturbances*, are an equally prominent feature of ecological systems worldwide (Lake 2000; Wolkovich *et al*. 2014). For instance, gradually shifting thermal regimes facilitate the poleward expansion of novel biotic assemblages (Zarzyczny *et al*. 2024), while continued losses of sea-ice cover drive annual declines in the reproductive success of keystone Antarctic species (Fretwell 2024). Note the distinction here between ramp (gradual, directional change) and press (constant and sustained pressure) disturbances (Fig. 1). Beyond human-driven global change, ramp disturbances are also a natural part of ecological systems, with successional changes in resource availability associated with gradual shifts in individual performance (Lasky *et al*. 2014), population dynamics (Martínez-Ramos *et al*. 2021), and community composition (Cline & Zak 2015). As such, predicting and managing the responses of ecological systems to ongoing global change requires understanding how the accumulation of impacts imposed by ramp disturbances influences their resilience over time (Ozgul *et al*. 2023; Paniw *et al*. 2019).

**Figure 1.**
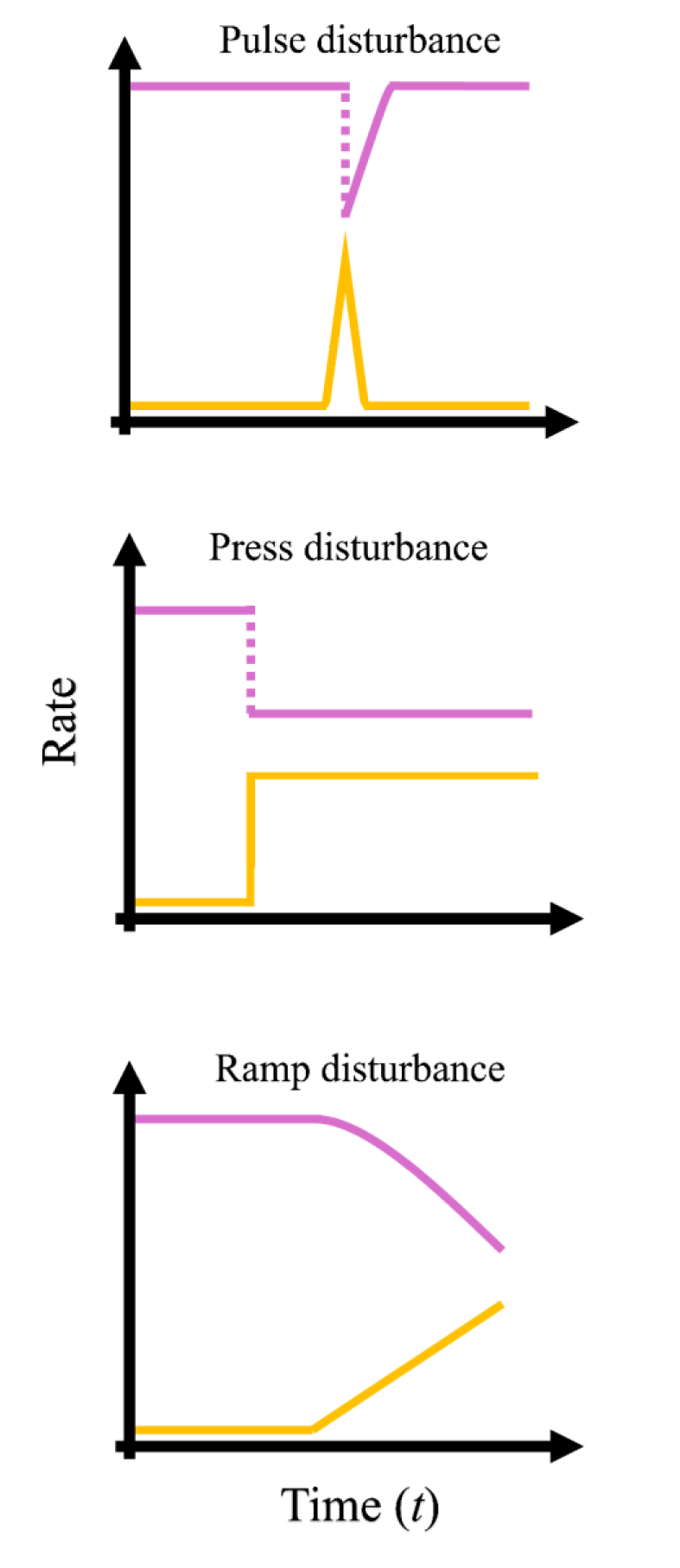
Ramp disturbances inflict very different perturbation regimes on the dynamics of natural populations, compared to pulse and press disturbances. The differing responses of population vital rates (pink, e.g. survival and fecundity) during their exposure to the disturbance intensity patterns (orange) associated with pulse-, press- and ramp disturbances.

Demographic resilience can be quantified by comparing short-term deviations in observed or forecasted population growth rates against stationary (*i.e.*, asymptotic) expectations (Capdevila *et al*. 2020). Under stationary conditions, in the absence of density feedback mechanisms, natural populations are expected to change in size at a constant rate over time, termed their asymptotic population growth rate (λ) (Caswell 2001; Groenendael *et al*. 1988). In reality, however, populations regularly experience periodic impacts to their survival and fecundity rates and/or their population structure (*i.e.,* the distribution of individuals across their life-cycle), leading to the expression of short-term (*i.e.*, transient) dynamics differing greatly from expected stationary trajectories (Koons *et al*. 2005; Stott *et al*. 2011). In this transient state, populations experience either increased (*i.e*., amplification) or decreased growth (*i.e.* attenuation) relative to asymptotic expectations. Quantifying a population’s potential for amplifying its instantaneous growth rate relative to λ reveals its capacity to benefit from a disturbance (Cant *et al*. 2023b; Jelbert *et al*. 2019). Meanwhile, quantifying a population’s attenuated growth rate relative to λ offers insight into its resistance potential (Capdevila *et al*. 2022b; Gamelon *et al*. 2014; McDonald *et al*. 2016; Stott *et al*. 2010). Combined, these metrics of amplification and attenuation constitute the demographic resilience framework (Fig. 2A) (Capdevila *et al*. 2020; Hodgson *et al*. 2015). Crucially, however, this framework centres upon the impacts of pulse disturbances (Capdevila *et al*. 2020). Thus, current predictions of population resilience overlook how the response of natural populations to disturbance events will be reshaped by the accumulation of gradual and directional vital rate shifts imposed by ramp disturbances.

**Figure 2.**
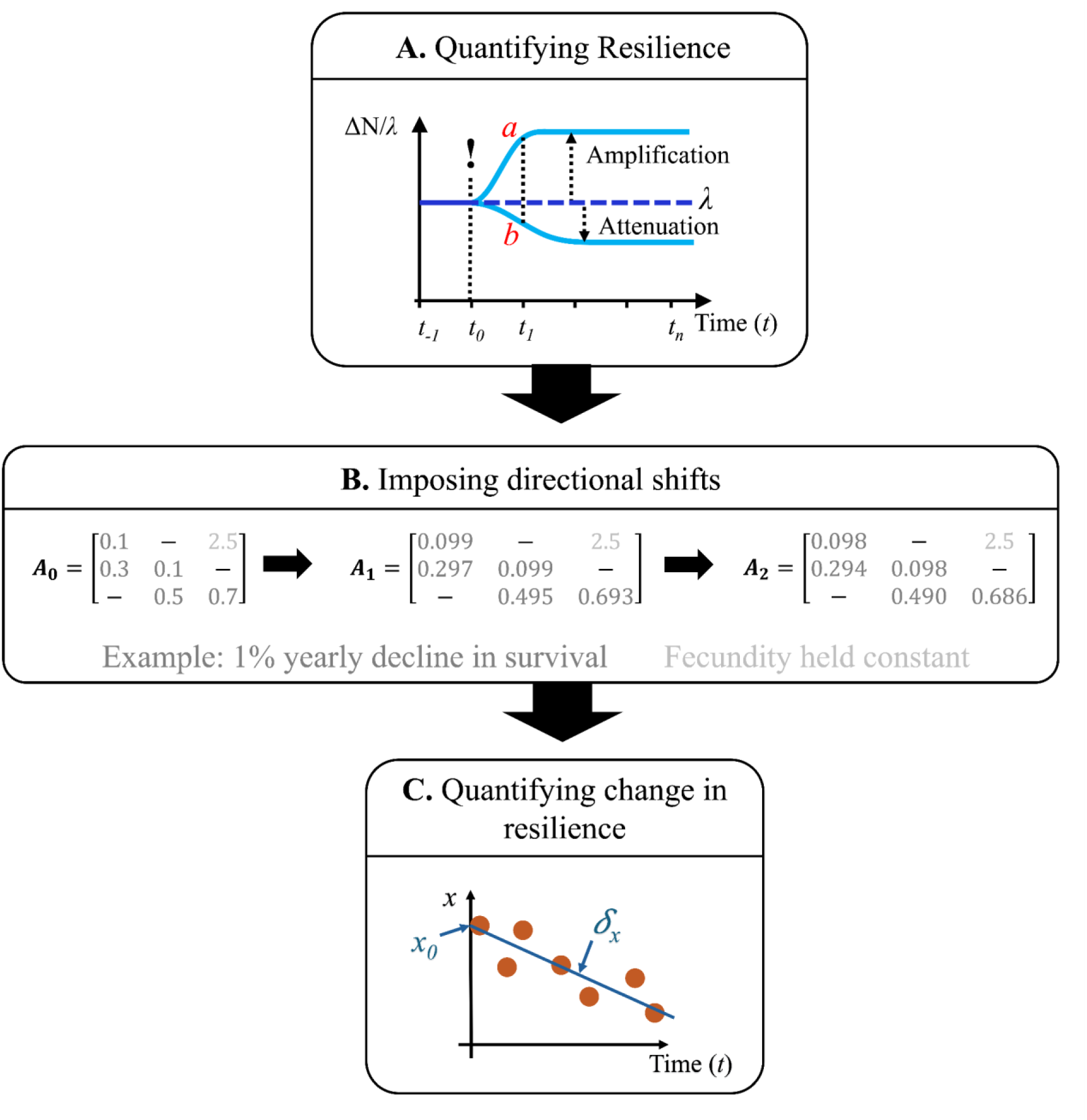
Pipeline used to evaluate how population resilience changes when exposed to ramp disturbances. **(A)** Under stationary conditions, populations are expected to change in size at a constant rate over time (i.e. their long-term population growth rate, λ). However, following a disturbance (!), populations can experience either amplification (*i.e.* increases in growth rate relative to λ) or attenuation (*i.e*. decreases in growth rate relative to λ). Quantifying these characteristics offers insight into population resilience. Here, we focused on estimates of amplification (*a*) and resistance (*b*). **(B)** We exposed a sample of matrix population models (MPMs) to a series of 50-year-long ramp disturbance scenarios. These scenarios entailed differing annual shifts in each population’s survival and fecundity (both independent and simultaneous shifts). **(C)** We calculated amplification and resistance estimates for each MPM along these time series, which we then used to estimate the rate at which the resilience of each population changed, per unit time, under each disturbance scenario *(S_x_*). Relating this rate of change back to established estimates of each population’s amplification and resistance (*x_0_*) then allowed us to assess how the change in resilience experienced by populations in response to ramp disturbances compares to our current perception of their demographic resilience following pulse disturbances.

Here, we simulate 50-year directional annual changes on the survival and fecundity of 511 populations across 344 species to examine how ramp disturbances could be expected to modify the resilience of natural populations (Fig. 2). Pulse and ramp disturbances inflict differing selective pressures on populations. For instance, some species promote their resilience to pulse disturbances through traits associated with structural integrity (*e.g.,* thicker bark [Pausas 2015], and robust morphologies [Darling *et al*. 2012]). Yet, these adaptations can leave individuals physiologically ill-equipped to handle gradual, long-term environmental shifts (Schoepf *et al*. 2019). Thus, we hypothesise that (H1) ramp disturbances will elicit greater magnitudes of change upon the resilience of populations that display higher amplification and/or resistance following pulse disturbances. We also predict that (H2) sessile species will experience greater magnitudes of change in their resilience, compared to mobile species. Sessile organisms often exhibit traits relating to slow population turnover (Reich 2014), dormancy/resprouting (Pausas & Keeley 2014), or structural resilience (Pausas 2015), all of which are beneficial for enduring pulse disturbances. In contrast, mobile species can simply move away to avoid discrete disturbances (Pausas 2019). Finally, (H3) we predict smaller shifts in population resilience in scenarios comprising opposite shifts in survival and fecundity, compared to those to which survival and fecundity respond in the same direction. This prediction corresponds with evidence that contrasting shifts in survival and fecundity can allow populations to maintain their growth rates (Doak & Morris 2010; Villellas *et al*. 2015), and that higher survival benefits resistance whilst enhanced fecundity promotes amplification (Cant *et al*. 2023a; Capdevila *et al*. 2022b).

## Materials & Methods

To assess the impacts of ramp disturbances on our current understanding of demographic resilience, we developed an analytical pipeline comprising the following steps: (1) Sampling structured population models, (2) Simulating directional environmental shifts, (3) Quantifying patterns of change in population resilience, and (4) Implementing comparative analyses.

### Sampling population models

To test the impact of ramp disturbances upon the resilience of natural populations, we extracted matrix population models (MPMs hereafter) for plant and animal species from the COMPADRE (version 6.23.5.0) (Salguero-Gómez *et al*. 2015) and COMADRE (version 4.23.3.1) (Salguero-Gómez *et al*. 2016) databases. COMPADRE and COMADRE contain MPMs describing the vital rates of survival, growth (progression & retrogression), and fecundity in discrete time along a life cycle made up of discrete stages for >12,000 populations across >1,200 species. Thus, to ensure comparability across our sample, we imposed a series of strict selection criteria. We only extracted MPMs consisting of three or more life stages, since lower dimension MPMs provide unreliable estimates of population transient dynamics (Koons *et al*. 2005). We only retained *mean* MPMs (*i.e.* the element-by-element mean MPM derived from a sequence of MPMs describing the dynamics of a single population through time) to mitigate against unrealistic transition probabilities arising from only observing a given population over only a single time period. In cases (24% of initial sample) where the same population was represented by multiple mean MPMs, we computed and retained the element-by-element mean of these multiple entries. This process entailed collapsing replicate MPMs to the dimensions of the smallest replicate (d) by collapsing together all life stages ≥ d into a single terminal stage using the *Rcompadre* R package (Jones *et al*. 2022); an approach that best maintains the eigenvalue structure of the original matrix (Salguero-Gómez & Plotkin 2010). Once we had a single *mean* MPM for each population, we collapsed all MPMs into 3 × 3 matrices.

Following initial MPM extraction, we filtered our sample based on the metadata provided with each MPM from the source study. To ensure the examination of demographic resilience in the wild, we focused on natural populations, thus excluding MPMs exposed to experimental treatments or parameterised with zoo/greenhouse data. To ensure our dataset represented population characteristics documented over consistent units of time, we retained only MPMs representing an annual survey periodicity. We also only retained MPMs determined to be ergodic, primitive, and irreducible, using the *popdemo* R package (Stott *et al*. 2012), ensuring we rejected MPMs comprising incomplete life cycles. Finally, we omitted MPMs for populations exhibiting clonality, due to the challenges associated with differentiating and comparing the dynamics of genets and ramets. These selection criteria resulted in an initial population sample of 561 populations (120 animals and 441 plants) from 372 species (Appendix S1). Note we did not account for pre- (*i.e.*, reproduction is a product of fecundity and offspring survival) *vs* post-reproductive (*i.e.*, reproduction is a product of adult survival and fecundity) MPMs. Post-reproductive MPMs are associated with higher values of recruitment (Jelbert *et al*. 2019). However, because the magnitudes of change we imposed upon each population’s dynamics during our ramp disturbance scenario were relative to their initial condition, the distinction between pre- and post-reproductive MPMs does not affect the compatibility of our measures of change in population resilience.

### Quantifying changes in population resilience & performance

To evaluate how the demographic resilience of natural populations to pulse disturbances changes following ramp disturbances, we imposed differing scenarios of changing survival and fecundity on each of our MPMs. These scenarios each correspond with a differing pairwise combination of percentage annual changes in the stage-specific profiles of survival and fecundity. The changes we imposed across each vital rate were: ±5%, ±4.5%, ±4%, ±3.5%, ±3%, ±2.5%, ±2%, ±1.5%, ±1%, ±0.7%, ±0.5%, ±0.3%, ±0.1%, & 0%, resulting in 729 differing pairwise combinations. To implement each scenario, we assigned each MPM obtained from COMPADRE or COMADRE as its population’s *initial condition* (***A****_0_*). We then exposed each population to the selected disturbance scenario over a 50-year period, quantifying the associated element-level changes across its MPM corresponding with the imposed changes to its state-specific survival probabilities and per-capita fecundity (***A****_t_*; Fig. 2B). Across our different ramp disturbance scenarios we halted simulations where survival and/or fecundity rates became biologically impossible (*i.e.*, survival probabilities < 0 or > 1 & fecundity rates < 0), retaining only outputs from iterations before the exceeding of biological limits. We also recognise that constant patterns of change in population vital rates likely overstates long-term impacts in systems with highly plastic responses and overlooks any non-linearities in population resilience introduced by density-dependence (Hernández *et al*. 2025). However, the absence of empirical data describing environment-driven ramp disturbances across species and the unique nature of density-dependent mechanisms across populations necessitate this approach.

To describe how measures of demographic resilience obtained for a given population change in response to ramp disturbances, we used the two indices: *^_Amplification_* and *^_Resistance_*. To obtain these measures, at each time step along our simulations of directional change described above, we used our sequence of ***A****_t_* products to calculate sequential estimates of the bounds of population reactivity *(p_i_;* maximum increase in population growth rate within one-time step of a disturbance, relative to stationary conditions) and first-step attenuation *(p_i_; ditto* maximum decrease). A proliferation of terms exists for describing different aspects of demographic resilience. Specifically, amplification is also referred to as demographic compensation (Capdevila *et al*. 2020), a term also used to refer to the process whereby opposing shifts in population vital rates allow for the maintenance of demographic performance (Doak & Morris 2010; Villellas *et al*. 2015). Thus, to distinguish between demographic processes and to make use of more common vernacular, we henceforth use amplification and resistance to refer to increases and decreases in population growth rates, relative to stationary conditions. We used the sequences of *p_i_* and *p_i_,* obtained for each population, to calculate our indices of change in demographic resilience under each scenario (Fig. 2C). Both *^_Amplification_* and *^_Resistance_* describe the rate at which the relative reactivity and first-step attenuation values of a population change, per unit time, during their exposure to ramp disturbances. We calculated these measures as the slope coefficients of linear regression models fit to the respective sequences of *p_i_* and *p_i_,* obtained for each population under each scenario. For each population, we also computed estimates of reactivity and first-step attenuation from their unmodified MPM (***A****_0_*). We retained these estimates as measures of our current perception of each population’s resilience to pulse disturbances, termed *Amplification_0_* and *Resistance_0_*, respectively.

To quantify the effect of our simulated ramp disturbance scenarios on population performance, we used the measure of stochastic population growth rate (*λs*). *λs* describes how variation across an iterative sequence of population growth rates, due to fluctuations in population vital rates over time, combines to inform a population’s long-term growth trajectory (Lewontin & Cohen 1969). For each population, we defined a starting population structure comprising 1000 individuals distributed according to the stable-state distribution (SSD) of the population’s initial MPM (***A****_0_*; equal to the dominant right eigenvector of ***A****_0_*). At each time step during our ramp disturbance simulations, we used our sequences of ***A****_t_* products to generate iterative estimates of population size (*N_t_*), calculating *λs* as the geometric mean of the sequence of *N_t_*/*N_t-i_* estimates produced for each population under each disturbance scenario.

Finally, we carried out data cleaning across our data sample. We omitted all estimates outside the 95% percentiles of each of our *^_Amplification_*, *^_Resistance_*, *Amplification*, and *Resistance_o_* variables. Next, we pruned our population sample to remove populations for which we had omitted estimates of *^_Amplification_*, *^_Resistance_*, *Amplification_o_* and *Resistance_o_* (14 populations), and for which we could not obtain phylogenetic details (see *Sourcing phylogenetic details*). This pruning resulted in a retained population sample of 511 populations (119 animals and 392 plants) comprising 344 species (Appendix S1). To ensure all variables were normally distributed, we then applied a power transformation to our estimates of *Amplification_0_* (−1 × *Amplification_0_*^-0.45^) and applied a cube root transformation to *S_Amplification_* (sgn(*x*) ^x^ VM)-

### Establishing phylogenetic relationships

To accommodate ancestral relatedness between populations of similar species, we computed the phylogenetic distances across our population sample. Keeping mobile and sessile species separate, we identified each species’ unique identifier within the Open Tree of Life (OpenTreeOfLife *et al*. 2019) (OTL; taxonomy version 3.6), which we used to construct species-level phylogenetic sub-trees using the *phytools* (Revell 2024) and *ape* R packages (Paradis & Schliep 2018). Briefly, we defined all plant and coral species as sessile. This definition resulted in a collection of populations composed largely of plant species (392 populations from 265 species) plus four populations from two coral species (*Paramuricea clavata* and *Pocillopora damicornis*). We classified all remaining animal species as mobile (115 populations from 77 species).

After computing branch lengths for our phylogenetic sub-trees (following Grafen’s computation method [Grafen 1989]) and ensuring each sub-tree was rooted (contains a single common ancestor) and clear of polytomies (≥ 3 species emerging from a single origin), we expanded the branch tips of species for which we had mean MPMs for multiple populations (15 animal species, and 51 plant species; Appendix S1). Implementing this expansion required replicate populations of the same species to be rooted onto the species’ corresponding branch tip, each separated by short branch lengths of infinitesimal size (ε = 0.0000001 units). This expansion approach assumes that populations of the same species are very closely related, whilst preventing polytomies. With recent work evidencing how analyses of demographic resilience are not sensitive to the order in which population replicates are introduced to population-level phylogenetic trees (Merrien *et al*. 2022), the order in which we added population replicates to our phylogenetic trees was random. During the construction of our population-level phylogenetic sub-trees, we dropped eight MPMs for which no phylogenetic information could be extracted from the OTL.

### Comparing change in resilience vs. established measures

To evaluate the relationship between established estimates of resilience to pulse disturbances (*Amplification_0_* & *Resistance_0_*) and the change in their resilience when exposed to ramp disturbances (*^_Amplification_* and *^_Resistance_*), we implemented a Bayesian multilevel modelling approach. Separately for our mobile and sessile groupings, we isolated estimates of *^_Amplification_* and *^_Resistance_* from scenarios involving changes in either survival or fecundity, during which the other vital rate remained unchanged. Individually, for scenarios involving changes in survival and fecundity, we then executed phylogenetically weighted regression models initiated with default priors using the *brms* R package (Bürkner 2017). In each case, we modelled the relationship between change in resilience following ramp disturbances (*^_Amplification_* or *^_Resistance_*) and established estimates of population resilience to pulse disturbances (*Amplification_0_* or *Resistance_0_*). We also included the fixed effect of disturbance scenario (*i.e.* the inflicted rate of change in either survival or fecundity), and a random-effect term allowing us to weight variance between populations corresponding to their phylogenetic distance. We fitted linear, polynomial and exponential model distributions, and used leave-one-out comparison (LOO) to select the best model fit in each case. Across all cases, the most appropriate model followed a polynomial distribution (Table S2). After selecting the most appropriate model fit, we ran our selected models over four Markov Chains, each consisting of 3,000 iterations, extracting the conditional effects of our independent variables on the observed patterns in *^_Amplification_* and *^_Resistance_*.

### Testing for evidence of compensatory mechanisms underlying population resilience

To assess whether populations can use contrasting shifts in their survival and fecundity rates to maintain their amplification and resistance, we again used a Bayesian multilevel modelling framework. Specifically, using outputs derived from our ramp disturbance simulations, we evaluated how changes in amplification (*^_Amplification_*), resistance (*^_Resistance_*), and stochastic population performance (*λs*) manifest across a two-dimensional parameter space describing the combinations of changes imposed by said ramp disturbances on survival *(^_Survival_*) and fecundity *(^_Fecundity_*). Again, our models were initiated using default priors and included a random effect term to account for phylogenetic variance-covariance. Likewise, we once again fitted our models using linear and polynomial distributions, before using LOO comparison to select the most appropriate distribution. Again, a polynomial distribution was deemed the most appropriate format in all cases (Table S3). We implemented this analysis using a Monte-Carlo resampling approach, each time selecting a 3% sample of our animal and plant datasets, running our selected model over a single Markov Chain consisting of 3,000 iterations. With this approach, for both our mobile and sessile groupings, we retained the mean predicted *^_Amplification_*, *^_Resistance_*, and *λs* values, estimated across 30 resampling iterations.

## Results

We found partial evidence that the magnitude of change populations experience in their resilience is greater in populations that, in response to pulse disturbances, possess higher amplification and/or resistance potential (H1) (Figs. 3 & 4). Across disturbance scenarios, only affecting either survival or fecundity, the magnitude of change in amplification (*^_Amplification_*) increased exponentially with increasing initial amplification estimates (*Amplification_0_*) (Fig. 3). Alternatively, populations exhibiting either minimal or high resistance in response to pulse disturbances (*Resistance_0_*) experience the lowest rates of change in their resistance under ramp disturbances (*^_Resistance_*) (Fig. 4). Across both amplification and resistance these patterns become more pronounced under increasing disturbance intensities. Note that declines in survival correspond with increases in *^_Amplification_* (Fig. 3A & B), whilst declines in fecundity result in increases in *^_Resistance_* (Fig. 4C & D). Accordingly, declines in either survival or fecundity displace a population’s dynamics such that the other rate becomes increasingly influential over time. Accordingly, the responses of *^_Resistance_* and *^_Amplification_* to shifts in survival and fecundity highlight how the relative balance between a population’s rates of survival and fecundity helps to determine its resilience, not just the observed changes in any one particular rate.

**Figure 3.**
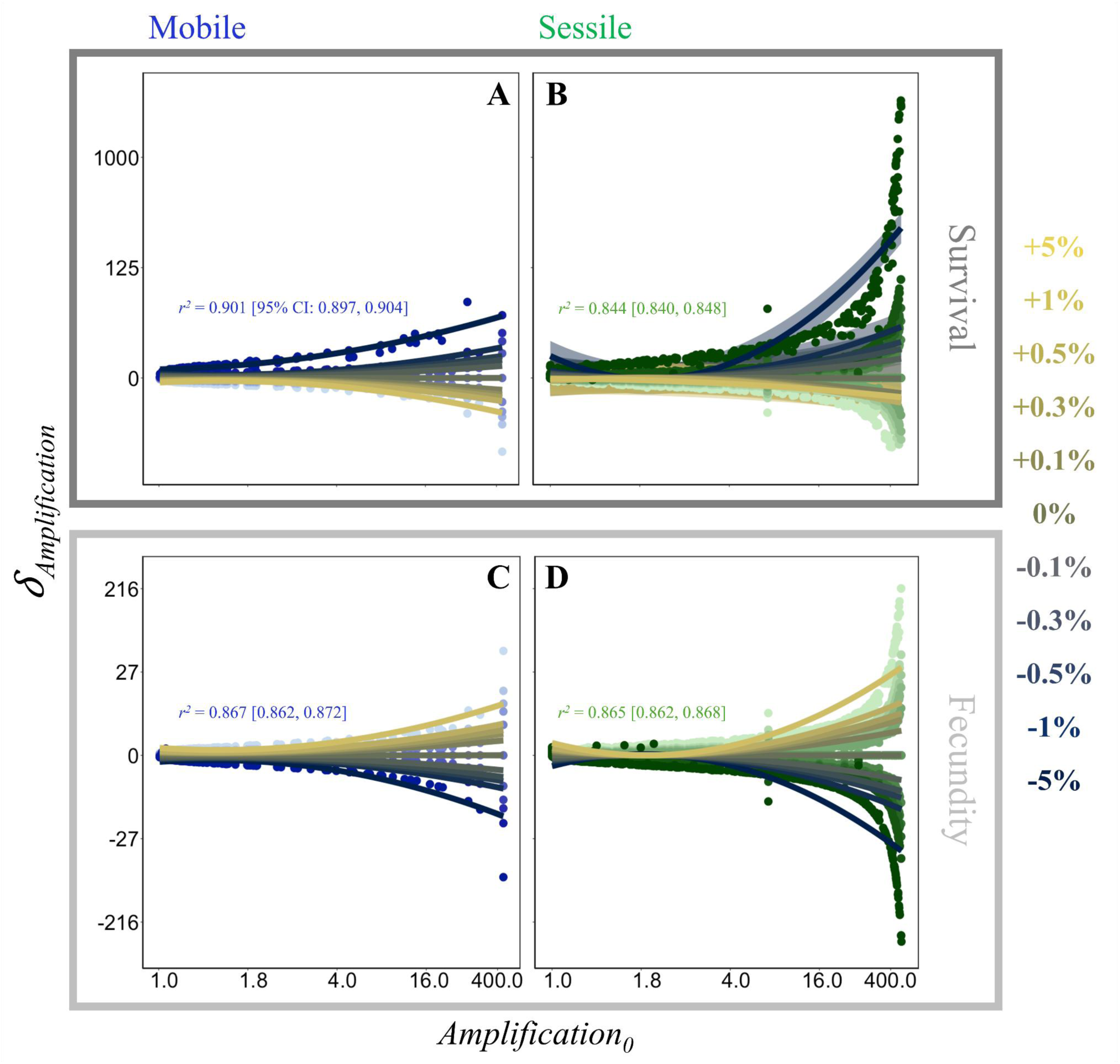
Populations expected to display greater amplification potential following pulse disturbances will experience a greater change in their amplification potential following exposure to ramp disturbances. The relationship between amplification to pulse disturbance (*Amplification_0_*) and the rate of change in amplification (*δ_Amplfication_*) when exposed to independent rates of directional change in survival and fecundity, in (**A & C**) mobile and (**B & D**) sessile populations (only patterns for ±5%, ±1%, ±0.5%, ±0.3%, ±0.1% & 0% change scenarios shown to ease visualisation). Solid lines represent the mean conditional effects extracted from phylogenetically weighted Bayesian multilevel regression models. Point colours are shaded based on their corresponding ramp disturbance scenario, ranging from the +5% rate of change scenario (lightest shading) to the -5% rate of change scenario (darkest shading). Error displayed as 95% CI.

**Figure 4.**
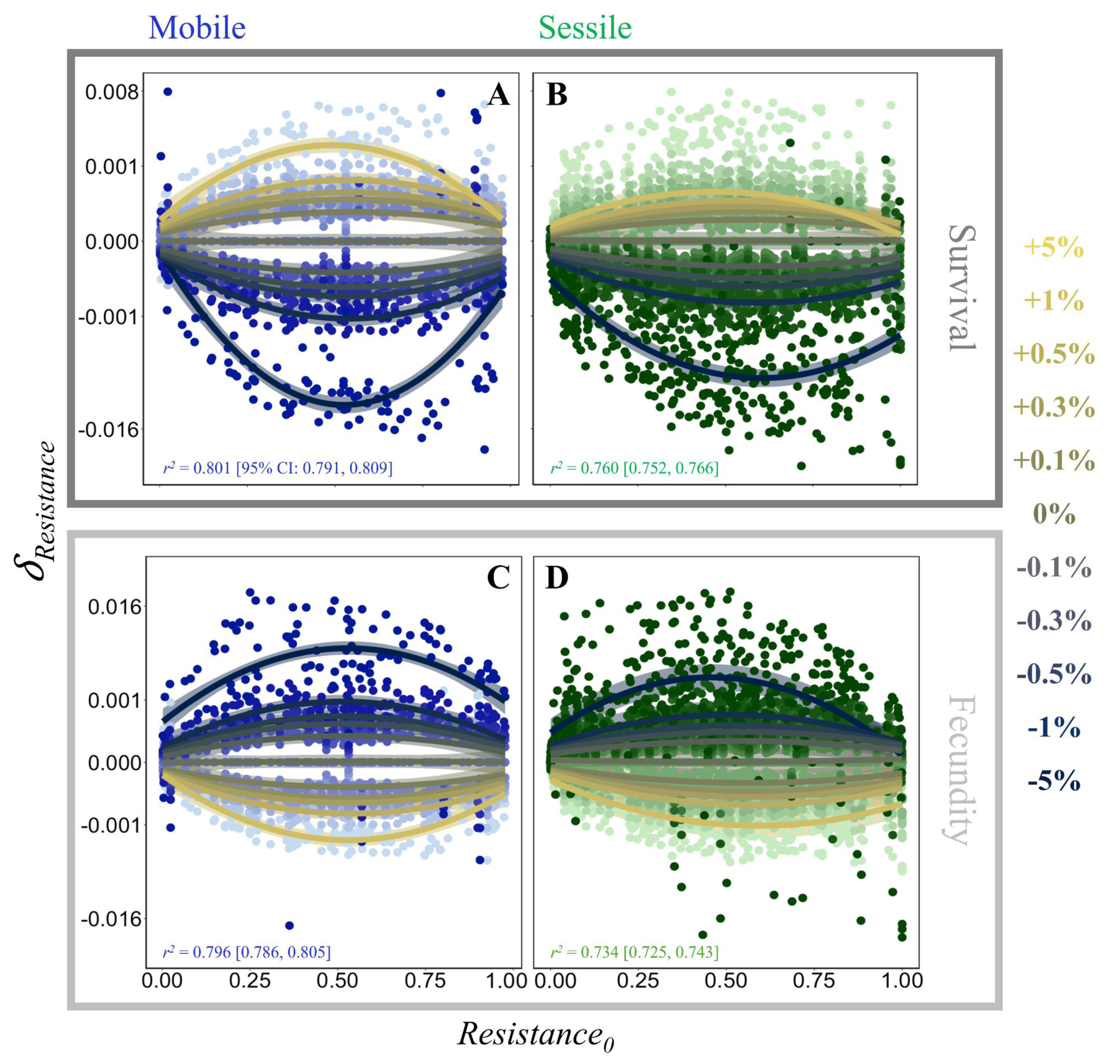
Populations expected to display either low or high resistance to pulse disturbances will experience less pronounced changes in their resistance following ramp disturbances. The relationship between resistance to pulse disturbance (*Resistance_0_*) and the rate of change in resistance (*δ_Resistance_*) when exposed to independent rates of directional change in survival and fecundity, in (**A & C**) mobile and (**B & D**) sessile populations (only patterns for ±5%, ±1%, ±0.5%, ±0.3%, ±0.1% & 0% change scenarios shown to ease visualisation). Solid lines represent the mean conditional effects extracted from phylogenetically weighted Bayesian multilevel regression models. Point colours are shaded based on their corresponding ramp disturbance scenario, ranging from the +5% rate of change scenario (lightest shading) to the -5% rate of change scenario (darkest shading). Error displayed as 95% CI.

Our findings partially indicate that changes in resilience following ramp disturbances are greatest in sessile species (H2). The relationship between change in amplification following ramp disturbances (*^_Amplification_*) and baseline amplification estimate (*Amplification*,) is most evident in sessile populations (Fig. 3). An annual 5% decline in survival elicits a rate of increase in amplification in sessile populations one order of magnitude greater than observed in mobile populations exhibiting comparable initial amplification (Fig 3A & B). Meanwhile, the relationship between *^_Resistance_* and *Resistance_o_* is more consistent across sessile and mobile populations (Fig. 4). However, this similarity does not mean that the relationship between *^_Resistance_* and *Resistance_o_* follows exactly the same shape across the two classifications. For instance, following an annual 5% decrease in survival, while mobile and sessile populations experience similar maximal declines in *^_Resistance_* (−0.0061 and -0.0104, respectively), the relationship tapers off more rapidly in sessile populations (Fig. 4A & B).

We found no evidence that contrasting shifts in survival and fecundity would counteract their individual effects on amplification and resistance (H3) (Fig 5). Demographic compensation occurs when populations maintain constant and viable long-term growth trajectories by offsetting declines in the rates of survival or fecundity with increases in the other vital rate (Doak & Morris 2010; Villellas *et al*. 2015). Indeed, demographic compensation is evident in *λs* across mobile and sessile species, with increases in survival compensating for the negative effects of declines in fecundity, allowing populations to maintain estimates of *λs* ≥ 1 (Fig. 5A & B). However, populations appear unable to counteract the effect of shifts in their vital rates on their resilience characteristics in the same way. Across both mobile and sessile populations, higher estimates of *^_Amplification_* are associated with scenarios of decreasing survival (lower *3_Survival_*) and increasing fecundity (higher *3_Fecundity_*) (Fig. 5C & D). Meanwhile, higher estimates of *3_Resistance_* are associated with scenarios of increasing survival and decreasing fecundity (Fig. 5E & F). Concurrently, if a ramp disturbance impacts the survival and fecundity of a given population at the same intensity and direction, the population will display no change in its estimated amplification, as indicated by the zero contours positioned diagonally across Fig. 5C & D. This observation is also true of resistance, although we report some evidence that increases in survival, at annual rates of >2%, can offset the negative influence of changing fecundity on population resistance *(^_Survival_* ^%^ 0.02 in Fig. 5E & F). Crucially, these mismatched patterns across population amplification, resistance, and *λs* warn that measures of population performance and resilience can express divergent responses to underlying vital rate shifts.

**Figure 5.**
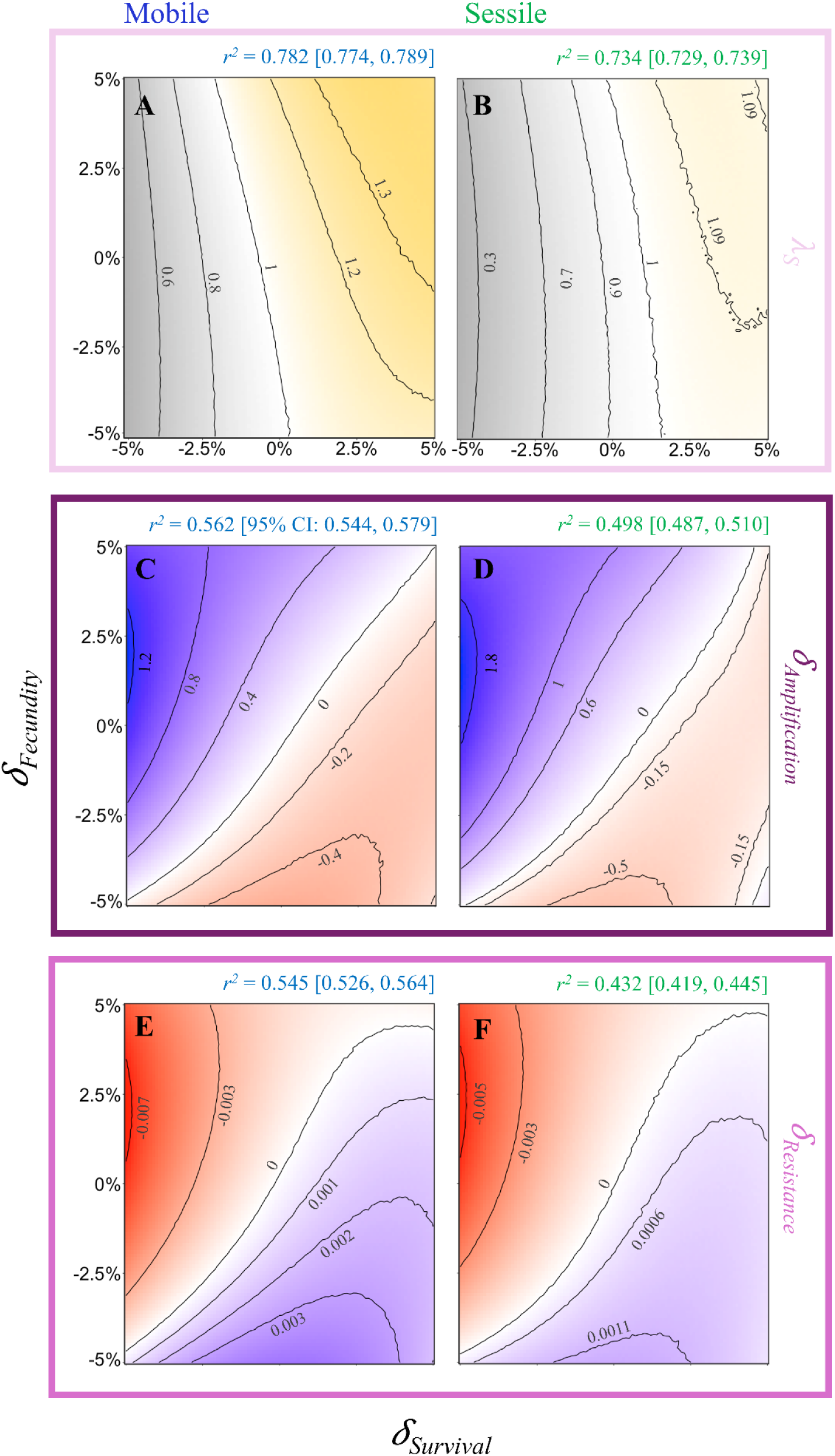
Patterns of change in population amplification and resistance associated with ramp disturbances may mask signals of population collapse. We separately quantified the relationships of changes in stochastic population growth rate (*λs*), amplification (*^_Amplification_*), and resistance (*^_Resistance_*), with simultaneous shifts in survival *(^_Survival_*) and fecundity *{^.._.uniy_*) across mobile (**A, C & E**) and sessile (**B, D & F**) populations. Colour gradients show mean predicted estimates of (*λs*), *3_Amplification_*, and *3_Resistance_*, extracted from 30 resampling iterations of phylogenetically weighted Bayesian multilevel regression models. Inset *r^2^* values indicate the overall strength of fitted models of the form *y ∼ ^_Survival_ * J_pecudnity_*, where *y* is the corresponding measure of (*λs*), *^_Amplification_*, or *^_Resistance_*. Dotted axis lines at 0% (*i.e.* no change) have been inserted across each panel to aid in the visualisation of changes in population resilience and performance across each parameter space.

## Discussion

Our findings illustrate how gradual, directional shifts associated with ramp disturbances could be expected to reshape and compound variation in the resilience of natural populations to pulse disturbances. Long-term population growth rates show contrasting responses to the respective ramp and pulse disturbance scenarios associated with changes in the mean and variability of climatic conditions (Morris *et al*. 2008). Logically, population resilience might also be expected to display divergent responses to the two disturbance types. To accommodate ramp disturbances within assessments of demographic resilience, we need to establish how the partitioning of energetic resources across survival and fecundity informs population resilience. The transient characteristics of natural populations offer insight into their resilience (Capdevila *et al*. 2020); a perception that represents the culmination of decades of effort to standardise assessments of population resilience (Hodgson *et al*. 2015; Ives *et al*. 2003; Neubert & Caswell 1997; Townley & Hodgson 2008). This framework has provided key insights into the evolution of resilience across taxonomic boundaries (Cant *et al*. 2023a; Guyennon *et al*. 2023; de Sousa *et al*. 2024; White *et al*. 2025), the drivers of vertebrate population declines (Capdevila *et al*. 2022a; Davis 2022), the geographic expansion of coral assemblages (Cant *et al*. 2023b), and the ecological implications of losing older individuals (Kopf *et al*. 2025). However, due to their nonlinear responses to the cumulative effects of ramp disturbances, existing metrics of demographic resilience cannot be used to forecast population resilience.

Against a backdrop of ongoing global change, before we can accurately forecast population resilience, we require a better appreciation for how population resilience is underpinned by the relative balance of survival *vs*. fecundity across the life cycle of organisms. By allowing populations to maintain longer-term growth rates (*i.e*., *λ* and *λs*) ≥ 1, demographic compensation underlies how species can adjust to localised climate shifts and helps to mediate range distributions (Doak & Morris 2010; Sheth & Angert 2018; Villellas *et al*. 2015). However, a similar mechanism appears absent in the context of population resilience, with populations unable to counteract the negative effects of changes in one vital rate on their amplification and resistance via changes to another. Moreover, provided the impacts of a ramp disturbance on the survival and fecundity of individuals within a population are of the same magnitude and direction, estimates of amplification and resistance will remain largely consistent while estimates of *λs* are expected to change. These divergent responses corroborate past evidence that, despite observed demographic characteristics being a manifestation of a population’s response to local environmental conditions (Lowe *et al*. 2017), exposure to environmental stochasticity does not select for particular resilience attributes (Cant *et al*. 2023a). Evidently, additional insight is needed to distil how, and whether, evolutionary selection pressures differ across characteristics of population resilience and demographic performance.

Investing in more enhanced transient dynamics, particularly higher amplification, is an advantageous strategy within more variable environments (Cant *et al*. 2023b; Ellis & Crone 2013; McDonald *et al*. 2016). Indeed, early successional plant species and mammals adopting a faster pace of life (*i.e.* shorter life spans, earlier maturity, and higher fecundity) exhibit a greater potential for demographic amplification (Gamelon *et al*. 2014; Stott *et al*. 2010), reflecting their reliance on boom-and-bust strategies for rapidly colonising available niche space. This mechanism parallels demographic lability, whereby large changes in some demographic parameter(s) allow populations to take advantage of good conditions (Ellis & Crone 2013; Hilde *et al*. 2020). However, our findings illustrate how fast-living species and populations, which possess greater amplification potential and/or a capacity for demographic lability, may be more sensitive to the negative impacts of ramp disturbances. This sensitivity could have cascading implications that would undermine the capacity for ecosystem recovery following physical disturbances. For instance, changing the composition of pioneer species assemblages shapes the post-fire recovery of forest communities (Gustafsson *et al*. 2021). Diminishing the viability of early successional species reliant on amplification, therefore, has the potential to inhibit future ecosystem recovery.

Our findings suggest that it may be difficult for interventions to enhance the resistance of species with already low resistance. There is a limit to the extent to which the dynamics of populations can maintain their viability under changing environmental scenarios (Rodríguez-Caro *et al*. 2021). Beyond this threshold, shifts in the demographic profiles of natural populations become increasingly indicative of population collapse (Cerini *et al*. 2023). We also show how species currently perceived as more resistant (higher *Resistance_0_*) may be better able to maintain their resistance when exposed to unfavourable shifts. However, it is important to consider that measures of resistance are bounded, and that the magnitude of population attenuation can only vary between 0 and 1. Subsequently, there is only so far one can increase the resistance of already highly resistant populations, and likewise, decrease the resistance of populations with already low resistance.

Intrinsic differences in the demographic profiles of populations associated with their relative mobility likely mediates their responses to ramp disturbances. Fundamentally, resistance relates to how vulnerable the most vulnerable individuals and/or systems are (Lawes *et al*. 2013; Pausas 2015). Therefore, although the mechanisms of their responses will be different, being resistant likely demands similar energetic investments across mobile and sessile species (Kooijman 2010). However, amplification relates to an increase in the number of individuals following a disturbance. Since sessile species cannot move away from disturbances, these species are expected to have evolved traits to maximise their fitness despite this constraint (*e.g.* resprouting & seed bank dormancy) (Lamont *et al*. 2011; Lennon *et al*. 2021). The physiological consequences of these traits will manifest quantitatively in the vital rates and structured population models developed for these species (*i.e.* high recruitment, presence of seed bank life stages & low seed to seedling survival) (Coulson *et al*. 2010). Thus, the greater response of amplification we report for sessile populations likely relates to their need to maintain their viability whilst having to endure environmental shifts. Alternatively, there is reduced need for mobile populations to amplify in response to environmental shifts as they can actively track their preferred conditions.

There are three key caveats that underlie our findings here. First, it is important to be vigilant in differentiating between true temporal shifts in resilience and mathematical artefacts. Calculating transient dynamics requires first normalising a population’s survival, development, and fecundity rates by its asymptotic growth rate to isolate transient and asymptotic behaviours (Townley & Hodgson 2008). However, during exposure to a ramp disturbance, as the survival and/or fecundity rates of a population change over time, it is necessary to account for the fact that the asymptotic growth rate value being used to normalise the dynamics of the population is not constant (Appendix S3). Second, our analyses focus on the transient bounds of population amplification and resistance, reflecting the most extreme possibilities in each metric given a population’s rates of survival, growth, and fecundity (Stott *et al*. 2011). Although, these bounds require no prior knowledge of population structure, the realised transient dynamics of any population depends not only on its survival, development, and fecundity rates, but also on the effect of a disturbance on its population structure (Koons *et al*. 2005). In the absence, however, of empirical insight quantifying the impacts of specific disturbance types on the dynamics of natural populations (Conde *et al*. 2019), our approach here allowed us to comprehensively explore the impacts arising from potential disturbance scenarios. Finally, while COMPADRE and COMADRE are a comprehensive source for demographic information on the world’s biota, they display the same geographic and taxonomic biases implicit across most ecological databases (Paniw *et al*. 2021; Römer *et al*. 2024; Salguero-Gómez *et al*. 2021). Whilst work is ongoing to remedy these biases (Salguero-Gómez *et al*. 2021), it is necessary that we consider how they inform our current perspectives.

We have illustrated how the exposure of natural populations to gradual and directional global change could be expected to alter our perception of demographic resilience. Our intention is to appraise previous efforts examining the resilience of populations to pulse disturbances, whilst cautioning against applying these tools to predict population resilience beyond the snapshot in time in which it was assessed (White *et al*. 2025). Indeed, efforts are ongoing to accommodate disturbances that maintain their intensity over time (*i.e.*, press disturbances) within our understanding of transient dynamics and demographic resilience (Tuljapurkar *et al*. 2023). In a field of research that is already rife with controversy and debate (Capdevila *et al*. 2021; Hodgson *et al*. 2015; Van Meerbeek *et al*. 2021), it is important to emphasise what existing frameworks can and cannot tell us when we consider the real-world complexities of natural systems (Hernández *et al*. 2025). Gradual, directional shifts in survival and fecundity associated with ramp disturbances will impose non-linear changes in the resilience of populations to pulse disturbances. Subsequently, predictions of population resilience using contemporary assessments of their resilience to pulse disturbances are not robust when set against a backdrop of ongoing and continuous global change. Instead, assessing population resilience over time requires quantifying the link between changes to a population’s relative investment across survival and fecundity and directional shifts in its amplification and resistance.

## Acknowledgements

We thank Prof. Tim Coulson and Dr Pol Capdevila for feedback on earlier versions of this work. J.C, C.H., and R.T. were supported by a NERC Pushing the Frontiers grant (NE/X013766/1) to R.S.G, I.S and A.H. C.H. was partly supported by a Marie Curie Fellowship (MSCA DensPopDy #10115386) with funding through UKRI (EP/Z002826/1), hosted by R.S.G.

## Data Availability Statement

All demographic data used in this study were extracted from the open-source COMPADRE and COMADRE databases (https://compadre-db.org/). Meanwhile, the phylogenetic distances between the species represented in the data sample used here were computed using the Open Tree of Life (https://tree.opentreeoflife.org); with the specific phylogenetic subtrees used in this study available for downloading from Zenodo (https://doi.org/10.5281/zenodo.18433775). The fully annotated R scripts used to extract and source this demographic and phylogenetic data are also publicly available on Zenodo via the same link.

## Code Availability Statement

The fully annotated R scripts used to carry out all the analyses presented in this manuscript are publicly available on Zenodo (https://doi.org/10.5281/zenodo.18433775).

