## Appendix for "Current estimates of population resilience are not robust to the longer-term impacts of directional environmental shifts"

**This PDF file includes:**

Supporting text

Figures S1 to S3

Tables S1 to S3

SI References

**S1: Details of species included in population sample.**

**Table S1. A list of the species included in the study.** The table shows the number of population replicates representing each species within the initially extracted population sample and the final data sample used in the study.

| Species | Populations (Initial Sample) | Populations (Final Sample) |
|---|---|---|
| *Acacia bilimekii* | 1 | 0 |
| *Actaea elata* | 1 | 1 |
| *Actaea spicata* | 2 | 2 |
| *Adenocarpus gibbsianus* | 1 | 1 |
| *Adenophora lobophylla* | 1 | 1 |
| *Adenophora potaninii* | 1 | 1 |
| *Aeschynomene virginica* | 1 | 1 |
| *Agrimonia eupatoria* | 1 | 1 |
| *Alces alces* | 3 | 3 |
| *Alliaria petiolata* | 12 | 12 |
| *Alnus incana subsp. rugosa* | 1 | 1 |
| *Alouatta seniculus* | 1 | 0 |
| *Amazona vittata* | 1 | 1 |
| *Ammocrypta pellucida* | 1 | 1 |
| *Anarrhinum fruticosum* | 1 | 1 |
| *Anemone patens* | 1 | 1 |
| *Anser anser* | 1 | 1 |

| Species | Populations (Initial Sample) | Populations (Final Sample) |
|---|---|---|
| *Anthropoides paradiseus* | 1 | 1 |
| *Anthyllis vulneraria* | 1 | 1 |
| *Antirrhinum lopesianum* | 1 | 1 |
| *Antirrhinum subbaeticum* | 2 | 2 |
| *Aquilegia chrysantha* | 1 | 1 |
| *Aquilegia sp.* | 1 | 0 |
| *Araucaria cunninghamii* | 1 | 0 |
| *Arenaria grandiflora subsp. bolosii* | 1 | 1 |
| *Argyroxiphium sandwicense* | 1 | 1 |
| *Arisaema serratum* | 1 | 1 |
| *Armeria maritima* | 1 | 1 |
| *Armeria merinoi* | 2 | 2 |
| *Artemisia genipi* | 1 | 1 |
| *Asclepias meadii* | 2 | 1 |
| *Aspasia principissa* | 1 | 1 |
| *Asplenium adulterinum* | 6 | 0 |
| *Asplenium cuneifolium* | 4 | 4 |
| *Asplenium scolopendrium* | 1 | 1 |
| *Astragalus alopecurus* | 1 | 1 |
| *Astragalus scaphoides* | 2 | 1 |
| *Astragalus tremolsianus* | 1 | 1 |
| *Astragalus tyghensis* | 4 | 4 |

| Species | Populations (Initial Sample) | Populations (Final Sample) |
|---|---|---|
| *Astroblepus ubidiai* | 1 | 1 |
| *Astrocaryum mexicanum* | 1 | 1 |
| *Astrophytum asterias* | 4 | 4 |
| *Astrophytum capricorne* | 2 | 2 |
| *Astrophytum ornatum* | 1 | 1 |
| *Atriplex acanthocarpa* | 1 | 1 |
| *Atriplex canescens* | 1 | 1 |
| *Avicennia germinans* | 1 | 1 |
| *Balsamorhiza sagittata* | 1 | 1 |
| *Banksia ericifolia* | 1 | 1 |
| *Bencomia exstipulata* | 1 | 1 |
| *Bertholletia excelsa* | 2 | 2 |
| *Boechera fecunda* | 3 | 3 |
| *Borassus aethiopum* | 1 | 1 |
| *Bostrychia hagedash* | 1 | 1 |
| *Brachyteles hypoxanthus* | 1 | 1 |
| *Brassica insularis* | 4 | 4 |
| *Broughtonia cubensis* | 1 | 1 |
| *Bursera glabrifolia* | 1 | 1 |
| *Buteo solitarius* | 1 | 1 |
| *Calathea ovandensis* | 1 | 1 |
| *Calidris temminckii* | 1 | 1 |
| *Callospermophilus lateralis* | 1 | 1 |
| *Calochortus lyallii* | 1 | 1 |
| *Calochortus macrocarpus* | 1 | 1 |
| *Calycophyllum spruceanum* | 1 | 1 |

| Species | Populations (Initial Sample) | Populations (Final Sample) |
|---|---|---|
| *Calyptorhynchus lathami* | 1 | 1 |
| *Canis lupus* | 1 | 1 |
| *Carapa guianensis* | 1 | 1 |
| *Carduus nutans* | 3 | 3 |
| *Carlina vulgaris* | 4 | 4 |
| *Carnegiea gigantea* | 1 | 0 |
| *Catopsis compacta* | 1 | 1 |
| *Cebus capucinus* | 1 | 1 |
| *Cecropia obtusifolia* | 1 | 0 |
| *Centaurea horrida* | 3 | 3 |
| *Cercopithecus mitis* | 1 | 1 |
| *Certhia americana* | 1 | 1 |
| *Cervus elaphus* | 2 | 2 |
| *Chamaecrista lineata var. keyensis* | 1 | 1 |
| *Chamaedorea radicalis* | 1 | 1 |
| *Cheirolophus metlesicsii* | 1 | 1 |
| *Chelodina expansa* | 1 | 1 |
| *Chelydra serpentina* | 1 | 1 |
| *Chen caerulescens* | 1 | 1 |
| *Chrysemys picta* | 1 | 1 |
| *Cirsium acaule* | 1 | 1 |
| *Cirsium palustre* | 1 | 1 |
| *Cirsium pannonicum* | 1 | 1 |

| Species | Populations (Initial Sample) | Populations (Final Sample) |
|---|---|---|
| *Cirsium perplexans* | 1 | 1 |
| *Cirsium pitcheri* | 5 | 5 |
| *Cirsium vulgare* | 1 | 1 |
| *Cleome droserifolia* | 1 | 0 |
| *Clidemia hirta* | 2 | 1 |
| *Coespeletia spicata* | 1 | 1 |
| *Coespeletia timotensis* | 1 | 1 |
| *Corallorhiza trifida* | 1 | 1 |
| *Cornus florida* | 1 | 1 |
| *Crocodylus johnsoni* | 3 | 3 |
| *Cryptantha flava* | 1 | 1 |
| *Cryptophis nigrescens* | 1 | 1 |
| *Cynoglossum officinale* | 1 | 1 |
| *Cypripedium calceolus* | 1 | 0 |
| *Cypripedium fasciculatum* | 3 | 3 |
| *Cyrtandra dentata* | 1 | 1 |
| *Cystoseira zosteroides* | 1 | 1 |
| *Cytisus scoparius* | 1 | 1 |
| *Dactylorhiza lapponica* | 2 | 2 |
| *Danthonia sericea* | 1 | 1 |
| *Daphne rodriguezii* | 5 | 5 |
| *Dasypus novemcinctus* | 1 | 1 |

| Species | Populations (Initial Sample) | Populations (Final Sample) |
|---|---|---|
| *Dendrophylax lindenii* | 1 | 1 |
| *Dicerandra frutescens* | 3 | 3 |
| *Dicorynia guianensis* | 1 | 1 |
| *Digitalis purpurea* | 2 | 0 |
| *Dioon caputoi* | 1 | 1 |
| *Dioon merolae* | 1 | 1 |
| *Dioon sonorense* | 1 | 1 |
| *Dioscorea chouardii* | 1 | 1 |
| *Dipsacus fullonum* | 1 | 1 |
| *Disporum smilacinum* | 1 | 0 |
| *Dracocephalum austriacum* | 5 | 5 |
| *Dypsis decaryi* | 1 | 1 |
| *Echinacea angustifolia* | 3 | 3 |
| *Echinocactus platyacanthus* | 2 | 2 |
| *Echinospartum ibericum subsp. algibicum* | 1 | 1 |
| *Eidolon helvum* | 1 | 1 |
| *Elephas maximus* | 1 | 1 |
| *Emydura macquarii* | 1 | 1 |
| *Encephalartos cycadifolius* | 1 | 1 |
| *Encephalartos villosus* | 1 | 1 |
| *Entandrophragma cylindricum* | 1 | 1 |
| *Eperua falcata* | 1 | 1 |

| Species | Populations (Initial Sample) | Populations (Final Sample) |
|---|---|---|
| *Epilobium latifolium* | 1 | 1 |
| *Epinephelus morio* | 1 | 1 |
| *Eriogonum longifolium var. gnaphalifolium* | 1 | 0 |
| *Eritrichium caucasicum* | 1 | 1 |
| *Erodium paularense* | 2 | 2 |
| *Eryngium alpinum* | 1 | 1 |
| *Eryngium cuneifolium* | 1 | 1 |
| *Eryngium maritimum* | 1 | 1 |
| *Erythronium japonicum* | 1 | 1 |
| *Escobaria robbinsorum* | 1 | 1 |
| *Escontria chiotilla* | 2 | 2 |
| *Eumetopias jubatus* | 1 | 1 |
| *Euphorbia fontqueriana* | 1 | 1 |
| *Euterpe edulis* | 1 | 1 |
| *Euterpe precatoria* | 1 | 1 |
| *Falco peregrinus* | 1 | 1 |
| *Fulmarus glacialis* | 1 | 1 |
| *Gardenia actinocarpa* | 1 | 1 |
| *Gasterosteus aculeatus* | 2 | 2 |
| *Gentiana pneumonanthe* | 1 | 0 |
| *Gentianella campestris* | 1 | 1 |

| Species | Populations (Initial Sample) | Populations (Final Sample) |
|---|---|---|
| *Geonoma macrostachys* | 1 | 0 |
| *Geonoma pohliana subsp. weddelliana* | 1 | 1 |
| *Geonoma schottiana* | 1 | 1 |
| *Geranium sylvaticum* | 1 | 1 |
| *Geum rivale* | 1 | 1 |
| *Giraffa camelopardalis* | 3 | 3 |
| *Gorilla beringei beringei* | 1 | 1 |
| *Gracilaria gracilis* | 2 | 1 |
| *Grias peruviana* | 1 | 1 |
| *Guarianthe aurantiaca* | 1 | 1 |
| *Gyps coprotheres* | 1 | 1 |
| *Haliaeetus albicilla* | 1 | 1 |
| *Helenium virginicum* | 1 | 1 |
| *Helianthemum juliae* | 1 | 1 |
| *Helianthemum polygonoides* | 1 | 1 |
| *Helianthemum teneriffae* | 1 | 1 |
| *Heliconia acuminata* | 1 | 1 |
| *Hilaria mutica* | 1 | 1 |
| *Himantoglossum hircinum* | 1 | 0 |
| *Hoplocephalus bungaroides* | 1 | 1 |
| *Horkelia congesta* | 1 | 1 |
| *Hyparrhenia diplandra* | 1 | 0 |

| Species | Populations (Initial Sample) | Populations (Final Sample) |
|---|---|---|
| *Hypericum cumulicola* | 1 | 1 |
| *Ipomopsis tenuituba* | 1 | 1 |
| *Iris germanica* | 1 | 1 |
| *Jacquiniella leucomelana* | 1 | 1 |
| *Jacquiniella teretifolia* | 1 | 1 |
| *Juniperus procera* | 1 | 1 |
| *Jurinea fontqueri* | 1 | 1 |
| *Khaya senegalensis* | 6 | 6 |
| *Kinosternon subrubrum* | 1 | 1 |
| *Kosteletzkya pentacarpos* | 1 | 1 |
| *Kummerowia striata* | 1 | 1 |
| *Lagopus muta* | 1 | 1 |
| *Lantana camara* | 2 | 2 |
| *Laserpitium longiradium* | 1 | 1 |
| *Lathyrus vernus* | 1 | 1 |
| *Lechea cernua* | 1 | 1 |
| *Lechea deckertii* | 1 | 1 |
| *Lepanthes acuminata* | 1 | 1 |
| *Lepanthes caritensis* | 1 | 1 |
| *Lepidium davisii* | 1 | 1 |
| *Liatris scariosa* | 1 | 1 |
| *Licania heteromorpha* | 1 | 0 |

| Species | Populations (Initial Sample) | Populations (Final Sample) |
|---|---|---|
| *Limonium carolinianum* | 1 | 0 |
| *Limonium delicatulum* | 1 | 0 |
| *Limonium erectum* | 1 | 1 |
| *Limonium malacitanum* | 1 | 1 |
| *Linum catharticum* | 1 | 1 |
| *Lomatium bradshawii* | 2 | 2 |
| *Lomatium cookii* | 1 | 1 |
| *Lophophora diffusa* | 1 | 1 |
| *Lotus arinagensis* | 1 | 1 |
| *Lupinus lepidus* | 1 | 1 |
| *Lupinus tidestromii* | 3 | 3 |
| *Lycaste aromatica* | 1 | 1 |
| *Macaca mulatta* | 1 | 1 |
| *Magnolia macrophylla var. dealbata* | 1 | 1 |
| *Malaclemys terrapin* | 1 | 1 |
| *Mammillaria crucigera* | 1 | 1 |
| *Mammillaria dixanthocentron* | 1 | 1 |
| *Mammillaria hernandezii* | 2 | 2 |
| *Mammillaria huitzilopochtli* | 1 | 1 |
| *Mammillaria magnimamma* | 1 | 1 |
| *Mammillaria napina* | 1 | 1 |
| *Mammillaria solisioides* | 1 | 1 |

| Species | Populations (Initial Sample) | Populations (Final Sample) |
|---|---|---|
| *Manilkara zapota* | 1 | 1 |
| *Marmota flaviventris* | 1 | 1 |
| *Mauritia flexuosa* | 1 | 1 |
| *Mezilaurus mahuba* | 1 | 1 |
| *Miconia albicans* | 1 | 1 |
| *Milvus migrans* | 1 | 1 |
| *Mimulus cardinalis* | 4 | 2 |
| *Mimulus lewisii* | 3 | 0 |
| *Minuartia obtusiloba* | 1 | 1 |
| *Mirounga leonina* | 1 | 1 |
| *Molinia caerulea* | 1 | 1 |
| *Mora paraensis* | 1 | 1 |
| *Nardostachys jatamansi* | 1 | 1 |
| *Neobuxbaumia macrocephala* | 1 | 1 |
| *Neobuxbaumia mezcalaensis* | 1 | 1 |
| *Neobuxbaumia polylopha* | 1 | 1 |
| *Neobuxbaumia tetetzo* | 1 | 1 |
| *Neotinea ustulata* | 5 | 5 |
| *Nipponia nippon* | 1 | 1 |
| *Nothofagus fusca* | 1 | 1 |
| *Odocoileus virginianus* | 2 | 2 |
| *Oenothera deltoides* | 1 | 1 |

| **Species** | **Populations (Initial Sample)** | **Populations (Final Sample)** |
|---|---|---|
| *Oncidium poikilostalix* | 1 | 1 |
| *Oncorhynchus clarkii* | 7 | 7 |
| *Oncorhynchus tshawytscha* | 2 | 2 |
| *Onychogalea fraenata* | 1 | 1 |
| *Orchis purpurea* | 1 | 1 |
| *Ovis canadensis* | 11 | 11 |
| *Oxandra asbeckii* | 1 | 1 |
| *Oxytropis jabalambrensis* | 1 | 1 |
| *Pachycereus pecten-aboriginum* | 1 | 1 |
| *Pan troglodytes schweinfurthii* | 1 | 1 |
| *Panax quinquefolius* | 2 | 2 |
| *Panthera pardus* | 1 | 1 |
| *Papio cynocephalus* | 1 | 1 |
| *Paramuricea clavata* | 3 | 3 |
| *Parkinsonia aculeata* | 1 | 1 |
| *Parolinia glabriuscula* | 1 | 1 |
| *Paronychia pulvinata* | 1 | 1 |
| *Pentaclethra macroloba* | 1 | 1 |
| *Periandra mediterranea* | 1 | 0 |
| *Persoonia bargoensis* | 1 | 1 |
| *Persoonia glaucescens* | 1 | 1 |
| *Petrophile pulchella* | 1 | 1 |
| *Phoebastria immutabilis* | 1 | 1 |

| Species | Populations (Initial Sample) | Populations (Final Sample) |
|---|---|---|
| *Phrynosoma cornutum* | 2 | 2 |
| *Phyllanthus emblica* | 2 | 2 |
| *Phyllanthus indofischeri* | 1 | 1 |
| *Pilosella floribunda* | 1 | 0 |
| *Pinguicula alpina* | 1 | 1 |
| *Pinguicula villosa* | 1 | 1 |
| *Pinus lambertiana* | 1 | 1 |
| *Pinus maximartinezii* | 1 | 1 |
| *Pinus nigra* | 1 | 1 |
| *Pinus strobus* | 1 | 1 |
| *Piriqueta cistoides subsp. caroliniana* | 1 | 1 |
| *Plantago coronopus* | 11 | 11 |
| *Platymiscium filipes* | 1 | 1 |
| *Pocillopora damicornis* | 1 | 1 |
| *Podocnemis expansa* | 1 | 1 |
| *Polemonium van-bruntiae* | 1 | 1 |
| *Polygonella basiramia* | 1 | 1 |
| *Primula elatior* | 1 | 1 |
| *Primula veris* | 1 | 1 |
| *Primula vulgaris* | 4 | 4 |
| *Propithecus edwardsi* | 1 | 1 |
| *Propithecus verreauxi* | 1 | 1 |

| Species | Populations (Initial Sample) | Populations (Final Sample) |
|---|---|---|
| *Prosopis glandulosa* | 1 | 0 |
| *Prosopis laevigata* | 1 | 1 |
| *Prunus africana* | 1 | 1 |
| *Prunus serotina* | 1 | 1 |
| *Pseudomitrocereus fulviceps* | 1 | 0 |
| *Pseudophoenix sargentii* | 3 | 3 |
| *Pterocereus gaumeri* | 2 | 1 |
| *Ptychosperma macarthurii* | 1 | 1 |
| *Purshia subintegra* | 1 | 0 |
| *Pyrrocoma radiata* | 5 | 5 |
| *Quercus rugosa* | 1 | 1 |
| *Ramonda myconi* | 2 | 2 |
| *Rangifer tarandus* | 1 | 1 |
| *Ranunculus peltatus* | 1 | 1 |
| *Rhizophora mangle* | 1 | 1 |
| *Rhopalostylis sapida* | 1 | 0 |
| *Rosmarinus tomentosus* | 3 | 0 |
| *Rumex rupestris* | 1 | 1 |
| *Sabal minor* | 1 | 1 |
| *Sabal yapa* | 1 | 1 |
| *Santolina melidensis* | 1 | 1 |
| *Sapium sebiferum* | 1 | 1 |

| Species | Populations (Initial Sample) | Populations (Final Sample) |
|---|---|---|
| *Saponaria bellidifolia* | 4 | 2 |
| *Sarcocapnos baetica* | 1 | 1 |
| *Sarcocapnos pulcherrima* | 2 | 1 |
| *Sarracenia purpurea* | 3 | 3 |
| *Saussurea medusa* | 1 | 1 |
| *Saxifraga aizoides* | 1 | 0 |
| *Scabiosa columbaria* | 1 | 1 |
| *Sceloporus grammicus* | 7 | 7 |
| *Scolytus ventralis* | 1 | 1 |
| *Scorzonera hispanica* | 1 | 1 |
| *Shorea leprosula* | 1 | 1 |
| *Silene acaulis* | 3 | 3 |
| *Silene douglasii var. oraria* | 1 | 1 |
| *Silene spaldingii* | 1 | 1 |
| *Silene tatarica* | 1 | 1 |
| *Sonchus pustulatus* | 1 | 1 |
| *Stenocereus eruca* | 1 | 1 |
| *Sterna hirundo* | 1 | 1 |
| *Sternotherus odoratus* | 1 | 1 |
| *Sternula antillarum browni* | 1 | 1 |
| *Strix occidentalis* | 1 | 1 |
| *Stryphnodendron microstachyum* | 1 | 1 |

| **Species** | **Populations (Initial Sample)** | **Populations (Final Sample)** |
| --- | --- | --- |
| *Succisa pratensis* | 4 | 3 |
| *Swietenia macrophylla* | 1 | 1 |
| *Syzygium jambos* | 1 | 1 |
| *Tamiasciurus hudsonicus* | 1 | 1 |
| *Telipogon helleri* | 1 | 1 |
| *Thalassarche melanophris* | 1 | 1 |
| *Thymus vulgaris* | 1 | 1 |
| *Tillandsia juncea* | 1 | 1 |
| *Tillandsia multicaulis* | 1 | 1 |
| *Tillandsia punctulata* | 1 | 1 |
| *Tillandsia violacea* | 1 | 1 |
| *Tolumnia variegata* | 1 | 1 |
| *Trillium camschatcense* | 1 | 1 |
| *Trillium grandiflorum* | 10 | 8 |
| *Trillium ovatum* | 1 | 0 |
| *Trillium persistens* | 4 | 4 |
| *Trollius europaeus* | 5 | 4 |
| *Trollius laxus* | 1 | 1 |
| *Tsuga canadensis* | 1 | 0 |
| *Umbonium costatum* | 1 | 1 |
| *Urocitellus armatus* | 1 | 1 |
| *Urocyon littoralis* | 1 | 1 |

| Species | Populations (Initial Sample) | Populations (Final Sample) |
| --- | --- | --- |
| *Ursus americanus* | 3 | 3 |
| *Ursus maritimus* | 1 | 1 |
| *Vella pseudocytisus subsp. paui* | 4 | 4 |
| *Verbascum fontqueri* | 3 | 3 |
| *Verticordia staminosa* | 1 | 1 |
| *Viola elatior* | 2 | 2 |
| *Viola pumila* | 2 | 2 |
| *Viola sagittata var. ovata* | 1 | 1 |
| *Vipera aspis* | 1 | 1 |
| *Virola surinamensis* | 1 | 1 |
| *Vitaliana primuliflora* | 1 | 0 |
| *Vouacapoua americana* | 1 | 0 |
| *Vriesea sanguinolenta* | 3 | 3 |
| *Vulpes vulpes* | 1 | 1 |
| *Xenosaurus grandis* | 1 | 1 |
| *Xenosaurus platyceps* | 2 | 2 |
| *Zalophus californianus* | 3 | 3 |
| *Zamia inermis* | 1 | 1 |
| *Zea diploperennis* | 1 | 1 |
| *Ziziphus jujuba* | 1 | 1 |

**S2: Leave-one-out (LOO) model comparisons.**

**Table S2.** LOO weightings comparing the suitability of linear, polynomial, and exponential distributions applied to the modelled relationships between measures of population resilience to pulse disturbances and the change in these resilience metrics when mobile and sessile populations experience directional shifts in their vital rates of survival and fecundity. LOO weights concerning patterns between $\delta_{Amplification}$ and $Amplification_0$ are highlighted in blue, whilst LOO weights concerning patterns between $\delta_{Resistance}$ and $Resistance_0$ are highlighted in red.

| | | **Modelled distribution** | | |
|---|---|---|---|---|
| | | **Linear** | **Polynomial** | **Exponential** |
| **Mobile** | **Survival** | 0.015 | 0.898 | 0.087 |
| | **Fecundity** | <0.001 | 0.883 | 0.117 |
| **Sessile** | **Survival** | 0.042 | 0.958 | <0.001 |
| | **Fecundity** | 0.010 | 0.990 | <0.001 |
| **Mobile** | **Survival** | 0.108 | 0.892 | |
| | **Fecundity** | 0.062 | 0.938 | |
| **Sessile** | **Survival** | 0.142 | 0.858 | |
| | **Fecundity** | 0.120 | 0.880 | |

**Table S3.** LOO weightings comparing the suitability of linear and polynomial distributions applied to models assessing how patterns in $\delta_{Amplification}$, $\delta_{Resistance}$, and $\lambda_S$ align across a two-dimensional parameter space defined by changes in survival and fecundity rates in mobile and sessile populations.

| | | Modelled distribution | |
|---|---|---|---|
| | | **Linear** | **Polynomial** |
| **Mobile** | **Amplification** | 0.107 | 0.893 |
| | **Resistance** | 0.123 | 0.877 |
| | $\lambda_S$ | 0.070 | 0.930 |
| **Sessile** | **Amplification** | 0.063 | 0.937 |
| | **Resistance** | 0.129 | 0.871 |
| | $\lambda_S$ | 0.067 | 0.933 |

**S3: Comparing patterns in long-term population growth rate, amplification, and resistance.**

We simulated a 5% annual increase in fecundity rates across 10 representative populations over 50 years. These simulations allowed us to demonstrate the simultaneous changes populations experience in their reactivity ($\underline{\rho}_1$; Fig. S1), first step attenuation ($\underline{\rho}_1$; Fig. S2), and asymptotic population growth rates ($\lambda$, Fig. S3), following continual and directional shifts in their underlying vital rates (*e.g.*, survival, fecundity). The 10 randomly selected populations comprised five plant and five animal populations. We sourced the structured population models describing the population dynamics of these selected populations from the COMPADRE & COMADRE databases (Salguero-Gómez *et al.* 2015, 2016).

Quantifying patterns of demographic resilience requires extracting metrics of transient dynamics from structured population models (Capdevila *et al.* 2020; Stott *et al.* 2011). For this extraction, it is necessary to isolate a population's transient dynamics from its asymptotic behaviour (Koons *et al.* 2005; Stott *et al.* 2011). This isolation is carried out by normalising the transitions in survival, development, and fecundity by dividing each element by the population's corresponding asymptotic growth rate (Townley & Hodgson 2008). However, using the simulations carried out here, we illustrate that during exposure to ramp disturbances, as the population's vital rates shift over time, its asymptotic growth rate also changes (Caswell 2001). Thus, the asymptotic growth rate used to isolate a population's transient dynamics is not consistent over time. Indeed, here we can see how the rates of change in reactivity and first-step attenuation experienced by populations exposed to ramp disturbances (Figs. S1 & S2) are equivalent to the rate of change observed in their asymptotic population growth rate (Fig. S3). The challenge of distinguishing this mathematical artefact from true temporal shifts in reactivity and first-step attenuation makes it difficult to assess changes in the transient dynamics of populations over time.

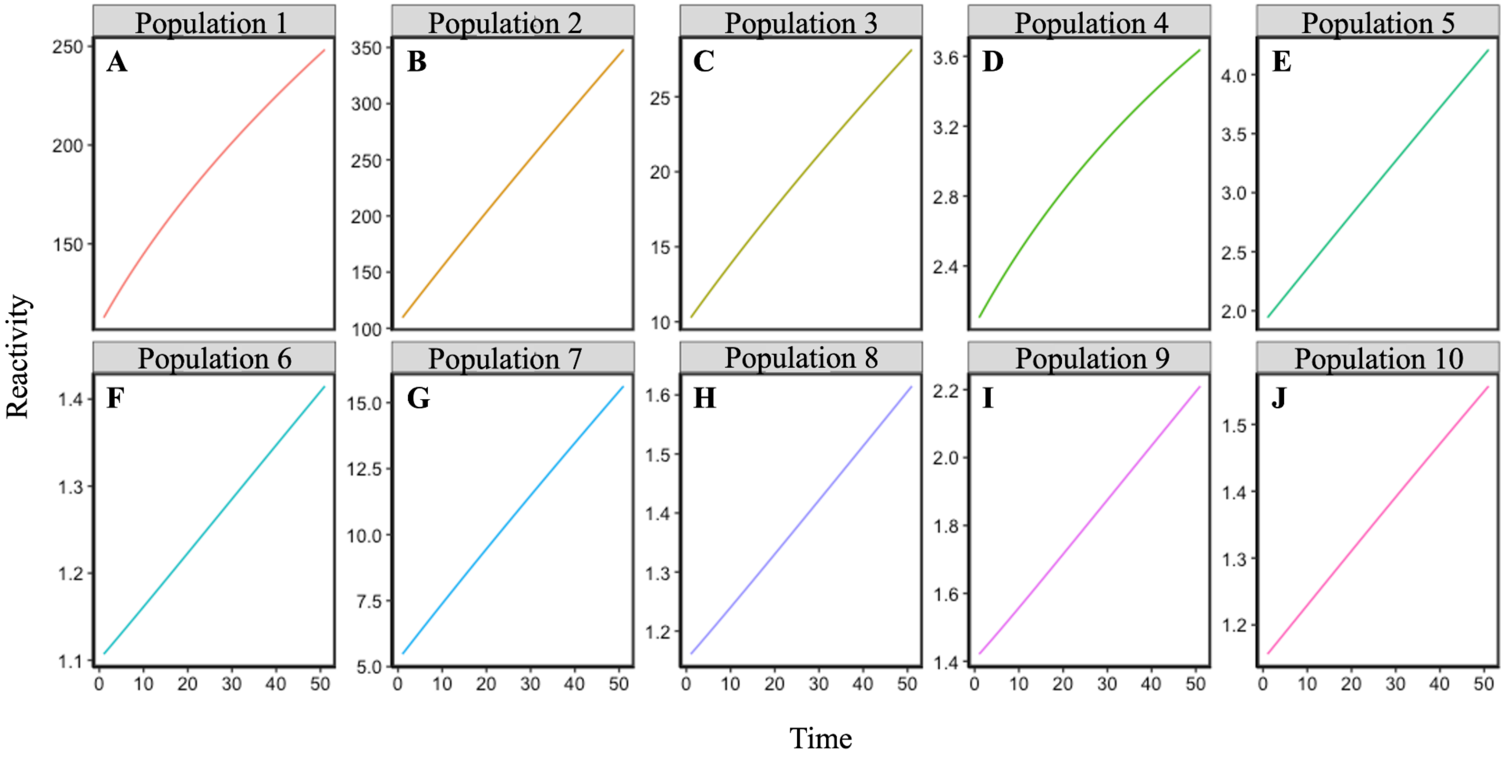


**Figure S1. Changes in population reactivity over time following exposure to a simulated ramp disturbance.** Iterative estimates of reactivity computed for 10 randomly selected plant **(A - E)** and animal **(F - J)** populations exposed to annual 5% increases in fecundity over a period of 50 years.

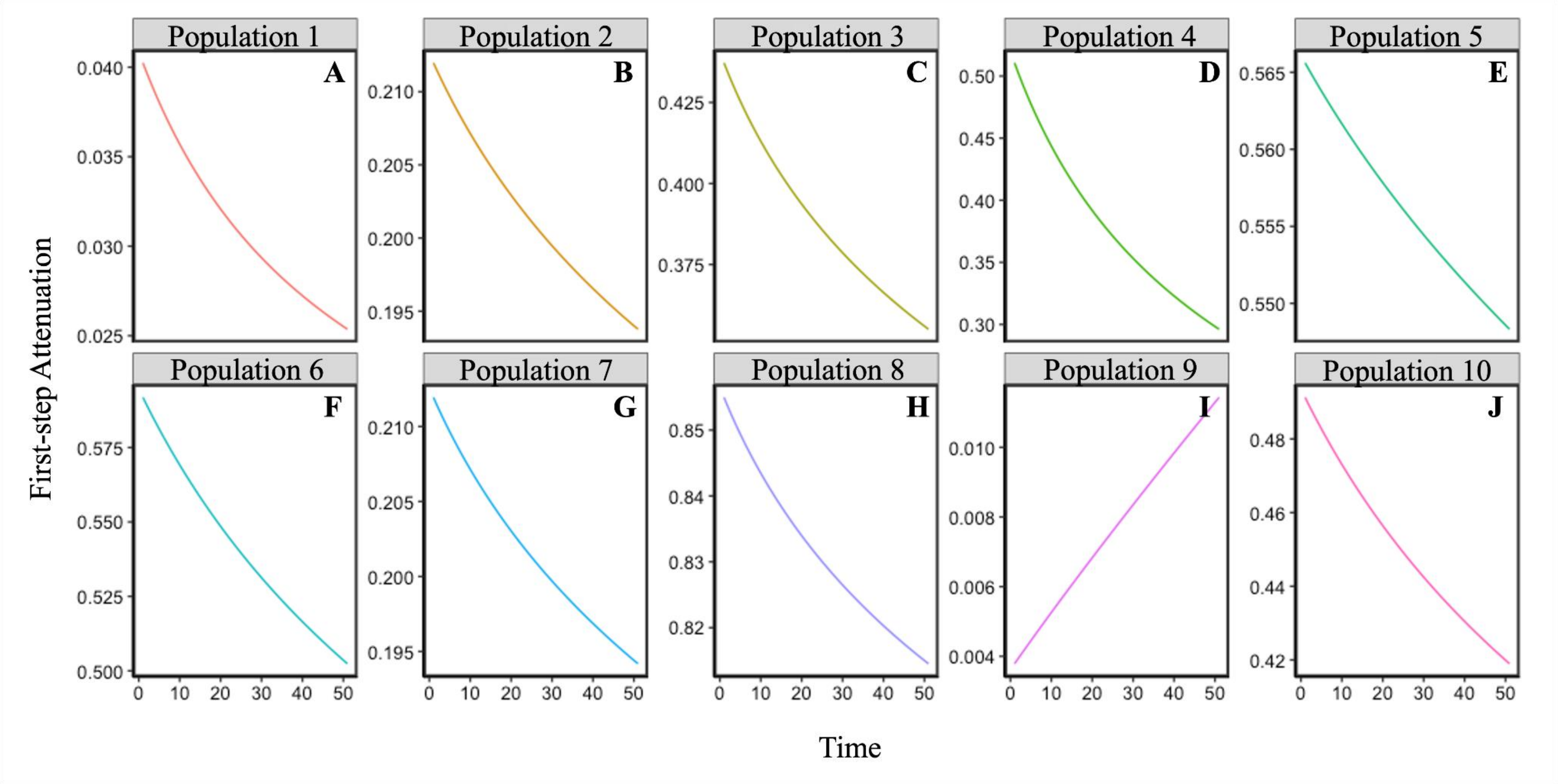


**Figure S2. Changes in first-step attenuation over time following exposure to a simulated ramp disturbance.** Iterative estimates of first-step attenuation computed for 10 randomly selected plant **(A - E)** and animal **(F - J)** populations exposed to annual 5% increases in fecundity over a period of 50 years.

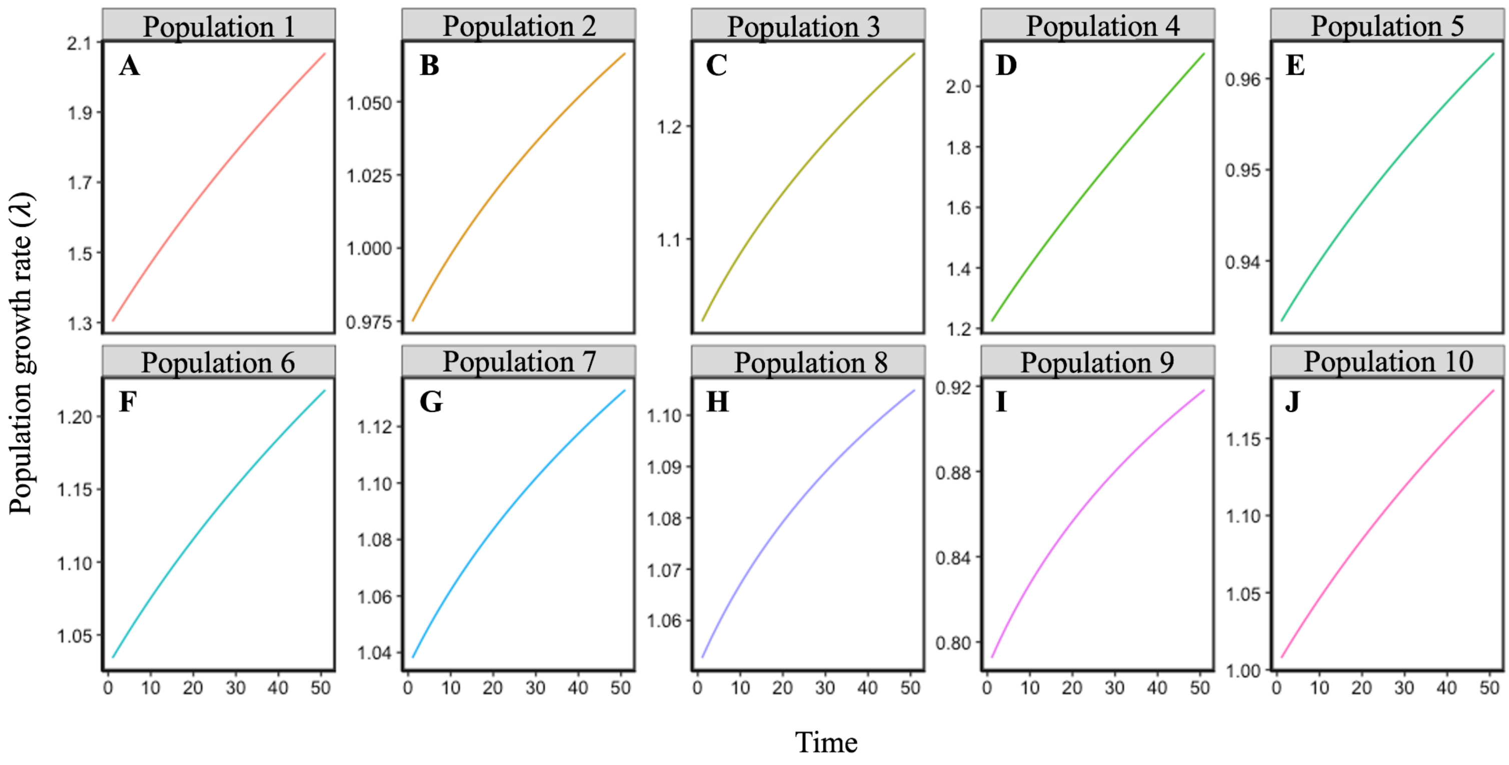


**Figure S3. Changes in asymptotic population growth rate (λ) over time following exposure to a simulated ramp disturbance.** Iterative estimates of first-step attenuation λ computed for 10 randomly selected plant **(A - E)** and animal **(F - J)** populations exposed to annual 5% increases in fecundity over a period of 50 years.
